# Recovering high-quality host genomes from gut metagenomic data through genotype imputation

**DOI:** 10.1101/2021.10.25.465664

**Authors:** Sofia Marcos, Melanie Parejo, Andone Estonba, Antton Alberdi

**Affiliations:** Applied Genomics and Bioinformatics, University of the Basque Country (UPV/EHU), Leioa, Bilbao, Spain; Center for Evolutionary Hologenomics, GLOBE Institute, University of Copenhagen, Copenhagen, Denmark

## Abstract

Metagenomic data sets of host-associated microbial communities often contain host DNA that is usually discarded because the amount of data is too low for accurate host genetic analyses. However, if a reference panel is available, genotype imputation can be employed to reconstruct host genotypes and maximise the use of such a priori useless data. We tested the performance of a two-step strategy to input genotypes from four types of reference panels, comprised of deeply sequenced chickens to low-depth host genome (~2x coverage) data recovered from metagenomic samples of chicken intestines. The target chicken population was formed by two broiler breeds and the four reference panels employed were (*i*) an internal panel formed by population-specific individuals, (*ii*) an external panel created from a public database, (*iii*) a combined panel of the previous two, and (*iv*) a diverse panel including more distant populations. Imputation accuracy was high for all tested panels (concordance >0.90), although samples with coverage under 0.28x consistently showed the lowest accuracies. The best imputation performance was achieved by the combined panel due to the high number of imputed variants, including low-frequency ones. However, common population genetics parameters measured to characterise the chicken populations, including observed heterozygosity, nucleotide diversity, pairwise distances and kinship, were only minimally affected by panel choice, with all four panels yielding suitable results for host population characterization and comparison. Likewise, genome scans between the two studied broiler breeds using imputed data with each panel consistently identified the same sweep regions. In conclusion, we show that the applied imputation strategy enables leveraging insofar discarded host DNA to get insights into the genetic structure of host populations, and in doing so, facilitate the implementation of hologenomic approaches that jointly analyse host genomic and microbial metagenomic data.

**Author summary:** We introduce and assess a methodological approach that enables recovering animal genomes from complex mixtures of metagenomic data, and thus expand the portfolio of analyses that can be conducted from samples such as faeces and gut contents. Metagenomic data sets of host-associated microbial communities often contain DNA of the host organism. The principal drawback to use this data for host genomic characterisation is the low percentage and quality of the host DNA. In order to leverage this data, we propose a two-step imputation method, to recover high-density of variants. We tested the pipeline in a chicken metagenomic dataset, validated imputation accuracy statistics, and studied common population genetics parameters to assess how these are affected by genotype imputation and choice of reference panel. Being able to analyse both domains from the same data set could considerably reduce sampling and laboratory efforts and resources, thereby yielding more sustainable practices for future studies that embrace a hologenomic approach that jointly analyses animal genomic and microbial metagenomic features.

## Introduction

The large molecular data sets generated through shotgun DNA sequencing usually contain useful information to characterise taxa, functions and structures beyond the primary aim of the study. This is especially true in metagenomic data sets that often present mixtures of DNA from eukaryotic, prokaryotic and viral origin (1,2). While primarily used for characterising the genomic architecture of microbial communities, metagenomic data generated from gut contents or faeces can also be used for extracting useful genomic information of the animal host (3). In fact, hologenomic approaches that entail joint analysis of animal genomes along with metagenomes of associated microorganisms to study animal-microbiota interactions, can benefit from such optimisation strategies (4,5).

However, mining host genomic data from metagenomic data sets entails a number of challenges. The fraction of host sequences in the metagenomic mixture is often unpredictable, and can range from a negligible proportion (<5%) to an almost complete representation (>95%) of the sample (6), even within a single taxon and sample type (7). Hence, a given amount of metagenomic sequencing effort does not ensure that the desired depth of host DNA sequencing will be reached. When the host DNA fraction in the metagenomic mixture is low, achieving the desired sequencing depth requires increasing sequencing effort, with its respective economic burden. In consequence, the amount of host DNA sequences generated is often insufficient for accurate variant calling.

One useful strategy for efficient data mining of host genomic information from metagenomic mixtures is genotype imputation, which consists in estimating missing haplotypes of poorly characterised genomes using a reference panel of high-quality genotypes (8). Using this approach, the information gaps of genomes with very low sequencing depth can be reconstructed based on the haplotype information of a properly characterised representative panel of genomes. Genotype imputation of single nucleotide polymorphisms (SNPs) is a widely employed approach in association studies to increase the density of variants of genomic data sets (9–11). In model organisms, the recent generation of large high-quality genomic databases, such as the human 1000 Genomes Project (12) and the 1000 Bull Genomes Project (13), has improved the accuracy of imputation and increased the statistical power of association analyses, especially for rare variants (14,15). However, ideal reference panels are only available for a limited number of model and farm species, and they also require high computational capacity.

When large reference panels are not available for small or isolated populations, an alternative strategy is to create a custom panel using a representative subset of genomes of the studied population (16,17). Due to its lower computational requirements, this approach can be more cost-efficient when studying closely related individuals, such as chickens from a given hatchery. This is because when haplotype diversity is limited, genomic information of a subset of the population can efficiently input haplotype information to the rest of the population. Moreover, the study-specific panel can be combined with individuals from public databases (16,17). This approach has been successfully employed in sheep (18), pig (19) and chicken (20) studies, for example.

Nevertheless, in addition to the size and diversity of the panel (21), imputation strategy may also affect the accuracy of recovered genotypes (22). In contrast to the standard imputation method, in which low density SNP arrays are imputed to high density based on a reference panel, shallow shotgun sequenced data displays particular challenges, as no genotype is known with certainty and SNPs may be distributed unevenly. Recently, a two-step imputation strategy for ultra low-depth coverage samples (<1x) was introduced (23). This approach relies on updating genotype likelihoods before imputing the missing genotypes using a reference panel in order to recover a higher density of SNPs with greater confidence. It was first proposed in human population genetics as an alternative to genotyping arrays for genome-wide association studies (23), and later applied to recover ancient human genomes (24). To the best of our knowledge, such a two-step imputation strategy has not been implemented yet in non-model animal populations with variable coverage and a limited number of available samples as a reference panel. Hence, there are no specific recommendations about the bioinformatic procedures for host genome recovery from metagenomic data sets and the choice of the most optimal panel to maximise accuracy of the imputation process. We also ignore how the choice of a custom reference panel could determine downstream analyses, such as measuring population genetics parameters.

Here, we present a straightforward approach to recover high-quality host genomes from gut metagenomic data, showcased in two broiler chicken breeds. We evaluate how the reference panel composition and sample depth of coverage affects imputation performance using four panels designed according to the resources scientists studying microbial metagenomics may have access to. We first calculate imputation accuracy between imputed and true genotypes in three chromosomes using 12 validation samples for which high-depth sequencing data is also available. Then, we employ a bigger data set of 100 individuals to impute all autosomal chromosomes and explore how the choice of the reference panel affects parameters commonly used in population genetics. Aiming at facilitating its implementation by other researchers, we provide the bioinformatic pipeline and guidelines for the choice of the most suitable panel and minimal depth threshold for a successful imputation.

## Methods

### Ethical statement

Animal experiments were performed at IRTA’s experimentation facilities in Tarragona under the permit FUE-2018-00813123 issued by the Government of Catalonia, in compliance with the Spanish Royal Decree on Animal Experimentation RD53/2013 and the European Union Directive 2010/63/EU about the protection of animals used in the experimentation.

### Target population and reference panels

Our study design involved genotype imputation from four reference panels with different origins and genetic features to a target chicken population characterised through low genomic coverage from intestinal metagenomic data.

### Target population

Genomic information of the target population of 100 chickens belonging to two broiler breeds (Ross 308 and Cobb 500, hereafter simply Ross and Cobb) was generated from metagenomic DNA extracted from the caecum contents of the birds. In short, ca. 100 mg of caecum content was collected right after euthanizing the animals and preserved in E-matrix tubes with DNA/RNA Shield buffer (Zymo Research, Cat. No. BioSite-R1200-125) at −20 °C until extraction. After physical cell disruption through bead-beating using a Tissuelyser II machine (Qiagen, Cat. No. 85300), DNA extraction was performed using a custom nucleic acid extraction protocol (details explained in (25)), and sequencing libraries were prepared using the adapter ligation-based BEST protocol (26). Paired-end 150 bp-long reads were generated on a MGISEQ-2000 sequencing platform over multiple sequencing lanes. Sequencing effort was decided based on the desired depth of the metagenomic fraction of the samples, which was the primary objective of the data generation. A preliminary screening revealed that caecum contents contain a large fraction of microbial DNA (>80-95%), and a limited relative amount of host DNA (< 5-15%) (Fig 1A). Aiming at about 15 GB (gigabases, ca. 50 million reads) of bacterial DNA per sample, caecum samples yielded between 0.5 and 4 GB of host DNA, which is equivalent to 0.5-4x depth of coverage of the chicken genome (~1.05 GB).

**Fig 1.**
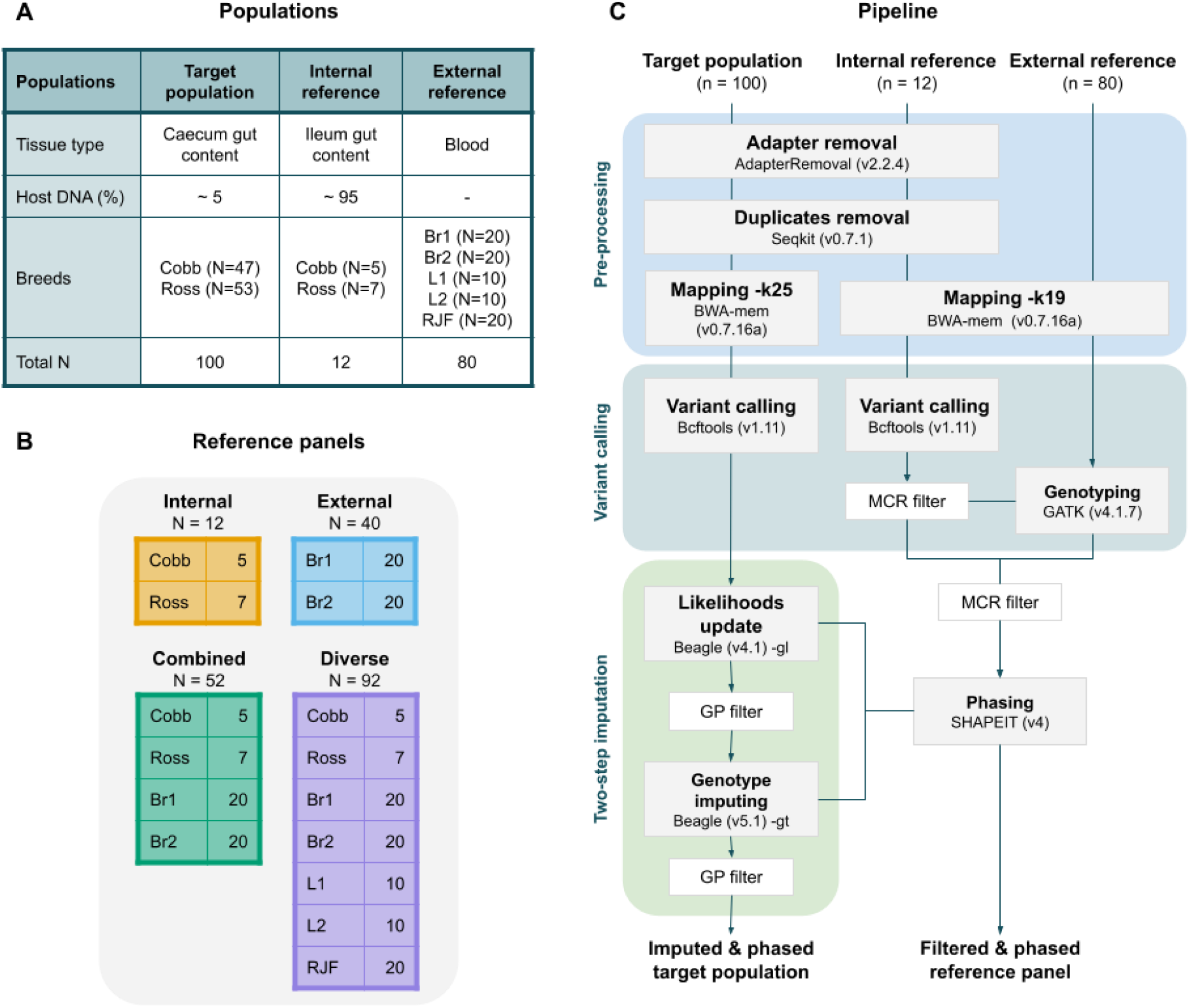
Study design and imputation pipeline for recovering host DNA. (A) The characteristics of the three data sets. (B) Composition and number of samples of the four reference panels used for imputation. Breeds are coded as Br1 = broiler line A, Br2 = broiler line B, L1 = white layer, L2 = brown layer, RJF= red junglefowl. (C) The study design has three data sets: the target population, internal reference and external reference samples. The bioinformatic procedure is divided into three steps: pre-processing, variant calling, and imputation. The input format of the starting step is a FASTQ file. After mapping we obtain a BAM file and from variant calling to the final step, procedures are performed using VCF file. The green box represents the steps proposed by Hui et al. (2020). Genotype probability (GP) filters are used during imputation and missing call rate (MCR) filters during panel design.

### Reference samples

Internal and external high-quality genome sequence data was used to create the reference panels. The internal reference data were generated from ileum content samples of 12 randomly selected individuals included in the target population (7 Ross and 5 Cobb), following the same procedures as explained above. In contrast to caecum samples, ileum contents contain a very large fraction (>90-95%) of host DNA, and a small representation of microbial DNA. Hence, in order to generate a comparable amount of microbial data to that of the caecum, ileum samples were sequenced aiming 100 GB/sample. This sequencing effort yielded about 90 GB of host DNA (ca. 80-90x depth of chicken genome), which enabled generating a high-quality internal reference panel from a subset of the studied population. In addition, chicken DNA sequence data of 40 broilers, 20 layers and 20 red junglefowls (RJF) generated by Qanbari et al. (2019) from blood samples were used as external reference data (Fig 1A).

### Composition of reference panels

We used different combinations of the internal and external reference samples to create the four reference panels used to evaluate imputation accuracy and impute the target population: *(i)* The internal panel comprised 12 animals from our target population (7 Ross and 5 Cobb), *(ii)* the external panel comprised 40 animals from two broiler breeds (different to our target population), *(iii)* the combined panel combined the previous two panels, and *(iv)* the diverse panel contained more distant populations (Fig 1B). The four panels varied in size and genomic diversity in order to see whether the composition of the reference panels affected imputation accuracy. With the internal panel, we tested if a small subset of the target population was enough for a proper imputation in low-quality host sequence data derived from metagenomic samples. The use of an external panel only was considered to test if it was a viable option for studies with a shortage of samples or a limited budget for high-depth host sequencing. The combined panel, on the other hand, permits combining both resources, the study-specific and database samples. Lastly, the diverse panel enabled us to test whether including distantly related individuals would be more effective than the three previously mentioned strategies.

### Pipeline for recovering host genotypes from metagenomic data

#### Data pre-processing

All the metagenomic sequence data we generated, which contained both host and microbial DNA, were pre-processed using identical bioinformatic procedures. In short, sequencing adapters were removed using AdapterRemoval (v2.2.4) (27) and exact duplicates using seqkit rmdup (v0.7.1) (28) prior to the read-mapping. Read-alignment to the chicken reference genome (galGal6; NCBI Assembly accession GCF_000002315.6) was conducted with BWA-MEM (v0.7.16a) (29). We employed default parameters except for the minimum seed length (-k), which was increased to 25 in order to reduce the number of incorrectly aligned read pairs. We added the flag -M, which was used to mark shorter split hits as secondary mappings. Aligned reads were sorted and converted into sample-specific BAM files before filtering out the metagenomic fraction (unmapped) using SAMtools view (v1.11) (30) with “-b” and “-F12” flags. Mapping statistics including depth and breadth of coverage as well as percentage of mapped reads were calculated using SAMtools’ depth and flagstat functions.

Pure genomic data (with no microbial fraction) generated by others (31) was downloaded from the EMBL-EBI ENA database, and mapped to the same chicken reference genome using BWA-MEM with -k default value and -M flag.

### Variant calling and genotyping

Variants in the target population were called by chromosome with the mpileup utility of SAMtools using standard parameters (-C 50 -q 30 -Q 20). Variant calling was performed with “-m” and “-v” flags to allow variants to be called on all samples simultaneously. Raw variants were filtered using BCFtools (v. 1.11) (32) commands “-m2”, “-M2” and “-v snps” to keep only bi-allelic SNPs. Variants of the internal reference samples were called the same way, but additionally, low quality variants with a lower base quality than 30 (QUAL<30) and variants with a base depth higher than three times the average (DP<(AVG(DP)*3) were removed to ensure only highly reliable variants were retained.

Since we were solely interested in imputing variants present in our target population, the external reference samples were genotyped by defining variant sites detected in the internal reference samples. Genotyping was performed for all autosomal chromosomes with GATK (v4.1.7.0) (33) HaplotypeCaller using the “--min-base-quality-score 20”, “--standard-min-confidence-threshold-for-calling 30”, “--alleles” and “-L” parameters to obtain calls at all given positions, followed by GATK SelectVariants "--select-type-to-include SNP” to only include SNPs.

In preliminary analyses, we also called variants in the external reference panel in order to examine the overlap with the variants present in the internal reference samples. We used the same procedures explained above for GGA1. Genotyping based on the positions of the internal panel and variant calling from scratch were compared by using the 40 broilers from the external reference panel for GGA1 (Fig 1B). A similar number of variants had been obtained for the genotyped (2.5 M) and the variant called VCF files (2.7 M). Moreover, 28% of the variants from the 40 broilers were not present in the internal reference samples (Fig S2). Thus, we decided to genotype the rest of the samples to reduce possible bias through the high number of variants specific to the external reference for the imputation of our target population.

### Two-step imputation via genotype likelihood updates

We imputed genotypes from the four aforementioned reference panels to the target population using a two-step strategy. Prior to imputation, the reference panels were filtered by excluding variants with missing genotypes to remove any potential noise caused by inference errors, and subsequently phased using SHAPEIT (v4) (34).

Imputation was performed in two steps following Homberg et al. (2019) and Hui et al. (2020). First, genotype likelihoods were updated based on one of the reference panels using Beagle 4.1 (35). Beagle 4.1 accepts a probabilistic genotype input with “-gl” mode, and it only updates sites that are present in the input file. Second, missing genotypes in the input file were imputed using Beagle 5.1 with “-gt” mode using the same reference panel. Beagle 5.1 only accepts files with a genotype format field, like later versions than Beagle 4.1. Therefore, the latest version cannot be used for both steps. Format field genotype probabilities (GP) were generated in both steps in order to enrich confident genotypes. We required the highest GP to exceed a threshold of 0.99 after both steps using BCFtools +setGT plugin. The rest of the parameters were set to default. Both steps’ input and output files were in VCF format. The schematic steps detailed in methods can be found in Fig 1C and the scripts in the following link (https://github.com/SofiMarcos/Host-genome-recovery.git).

### Imputation accuracy using 12 validation samples

The accuracy of the imputation using the four reference panels was tested using the 12 individuals for which we generated both low-depth (target population) and high-depth (internal reference samples) sequence data from caecum and ileum contents, respectively, hereafter referred to as validation samples. The low-depth samples of the 12 individuals had a depth of coverage spanning 0.05x to 3.73x. For an unbiased evaluation, we employed a leave-one-out cross-validation (LOOCV) approach by excluding each of the 12 validation samples once from the reference panel in each of the different imputation scenarios. Considering the large size-variation of avian chromosomes, a macrochromosome (GGA1, 197.6 MB), a mid-size chromosome (GGA7, 36.7 MB) and a microchromosome (GGA20, 13.9 MB) were selected for the test to optimise runtime and computational resources. Concordance between the internal reference samples and imputed genotypes was calculated for each individual chicken using VCFtools, with the“--diff-discordance-matrix” option. Precision of heterozygous sites was also calculated, since these alleles are the most difficult to impute correctly. Kruskal-Wallis test was performed to test for differences across chromosomes. A paired sample T-test and F-test were performed for both parameters to verify if the difference in means and variances were significant between reference panels. T-test p-values were adjusted using Bonferrini’s correction method. Moreover, imputation accuracy was estimated for variants in different minor allele frequency (MAF) bins to evaluate whether rare and common variants are equally correctly imputed. We thus extracted variant frequencies from the internal panel by analysing precision of heterozygous (het.) sites for the GGA1 in bins of 0-0.05, 0.05-0.1, 0.1-0.3 and >0.3.

### Impact of reference panel on population genetics inference

We explored the implications of using different reference panels in downstream analyses of population genetic inferences, including population structure, genetic diversity, and genome scans for signatures of selection.

These analyses were run in all but two outlier samples with depths of coverage of 0.07x and 0.05x, which were below the threshold of 0.28x corresponding to the lowest successfully imputed sample in the validation set (genotype concordance of >0.90 and het. sites precision of >0.75, see results below). We thus used 100 samples (53 Ross and 47 Cobb) for which we ran the host DNA recovery pipeline for all the autosomal chromosomes and analysed common population genetics parameters including observed heterozygosity (O.Het), nucleotide diversity (π), pairwise distance as estimated through identity-by-state (1-IBS) and kinship. The same analyses were also conducted for 10 validation samples (for the low-depth and high-depth samples) after excluding two of them, whose respective counterparts in the target populations (with 0.05 and 0.07x depth) were filtered out. The imputed data sets with each of the panels were filtered for missingness 0 with PLINK (v1.9) (36).

For measuring population genetics parameters, the VCF files were filtered for MAF >0.05. O.Het was calculated for each individual using the command “--het” in PLINK (v1.9). π was calculated in 40 kb windows with 20 kb step size across autosomal chromosomes using VCFtools. For the validation samples whole-genome windowed values were averaged to generate a genome-wide π for each individual. For the target population, π was calculated for each breed population. Paired sample T-tests were performed for O.Het and π parameters. Pairwise distance was calculated using “--distance square 1-ibs” in PLINK (v1.9). Kinship was calculated with the command “--make-king square” using PLINK (v2). To test the correlation between the resulting matrices from the pairwise distance and kinship analyses using different panels, a Mantel test was performed with the R package ade4 (37).

We further tested whether genome scans for selection between the Cobb and Ross population with each of the imputed datasets yielded consistent results. To this end, we calculated population differentiation along the genome using fixation index (FST) between both breeds using each panel. FST was calculated in sliding windows of 40 kb with 20 kb overlap across autosomal chromosomes. Window-based FST values were then normalised, and regions with values above the 99th and 99.9th percentile were considered as putative selective sweep regions (38). The overlap of these regions across the datasets using the different reference panels were used as an estimate of consistency.

## Results

### Alignment and coverage

The mapping statistics of the 100 samples used to characterise the target population (caecum content) and the 12 internal reference samples (ileum content) were drastically different. Caecum samples showed an average of 1.84±2.35x (mean±SD) depth of coverage and 52.41±24.20% of breadth of coverage. Ileum samples had 92.70±7.64% of host DNA and an average depth of 93.16±9.07x, practically covering the entire reference genome (98.89±0.01%).

### Pipeline fitting

The pipeline required some tests and adjustments to optimise it to our system. The standard alignment (seed length 19) presented an unconventional distribution of reads across the genome, i.e. unspecified read mapping leading to regions being stacked with 80+ reads (Table S1). In order to remove as many remaining microbial reads as possible, we increased the seed length to 25. Standard deviation of the depth of coverage decreased considerably (from 202.79 to 3.66), while the mean depth decreased from 2.78x to 1.73x. The breadth of coverage decreased by 9% (Fig S1).

### Imputation accuracy of 12 validation samples

The internal (n=12), external (n=40), combined (n=52) and diverse (n=92) reference panels were used to study (i) the effect of panel size and diversity and (ii) sample depth of coverage threshold on imputation accuracy in three chromosomes with contrasting dimensions. Variant calling in the internal reference samples detected 2.4 M, 470 K and 182 K putative SNPs in chromosomes GGA1, GGA7 and GGA20, respectively. After genotyping the external reference samples and combining them to create the external, combined and diverse panels, each panel was filtered before being phased. As a consequence, the filtering step decreased the number of SNPs by 13.83±1.36% for the external and combined, and by 23.80±0.99% for the diverse panel, which yielded panels with different numbers of SNPs (Fig 2A). More than 96% of the total SNPs in each panel successfully passed the multiple filters of the pipeline, even for samples with less than 1x coverage (Fig 2B). Furthermore, the proportion of imputed SNPs increased and gained uniformity across samples when the panel was larger but had fewer SNPs. The mean number of imputed SNPs across samples differed between all the panels: internal vs external (t=14.58, p-value < 0.001), external vs combined (t=13.56, p-value < 0.001) and combined vs diverse (t=11.63, p-value < 0.001). The F-test to compare variances was significant only between the diverse and the rest of the panels: internal vs diverse (F= 30.54, p-value<0.001), external vs diverse (F=24.24, p-value< 0.001) and combined vs diverse (F= 11.31, p-value<0.001). Results indicate that the variance across samples for the diverse panel greatly decreased compared to the rest of the panels (Fig 2B).

**Fig 2.**
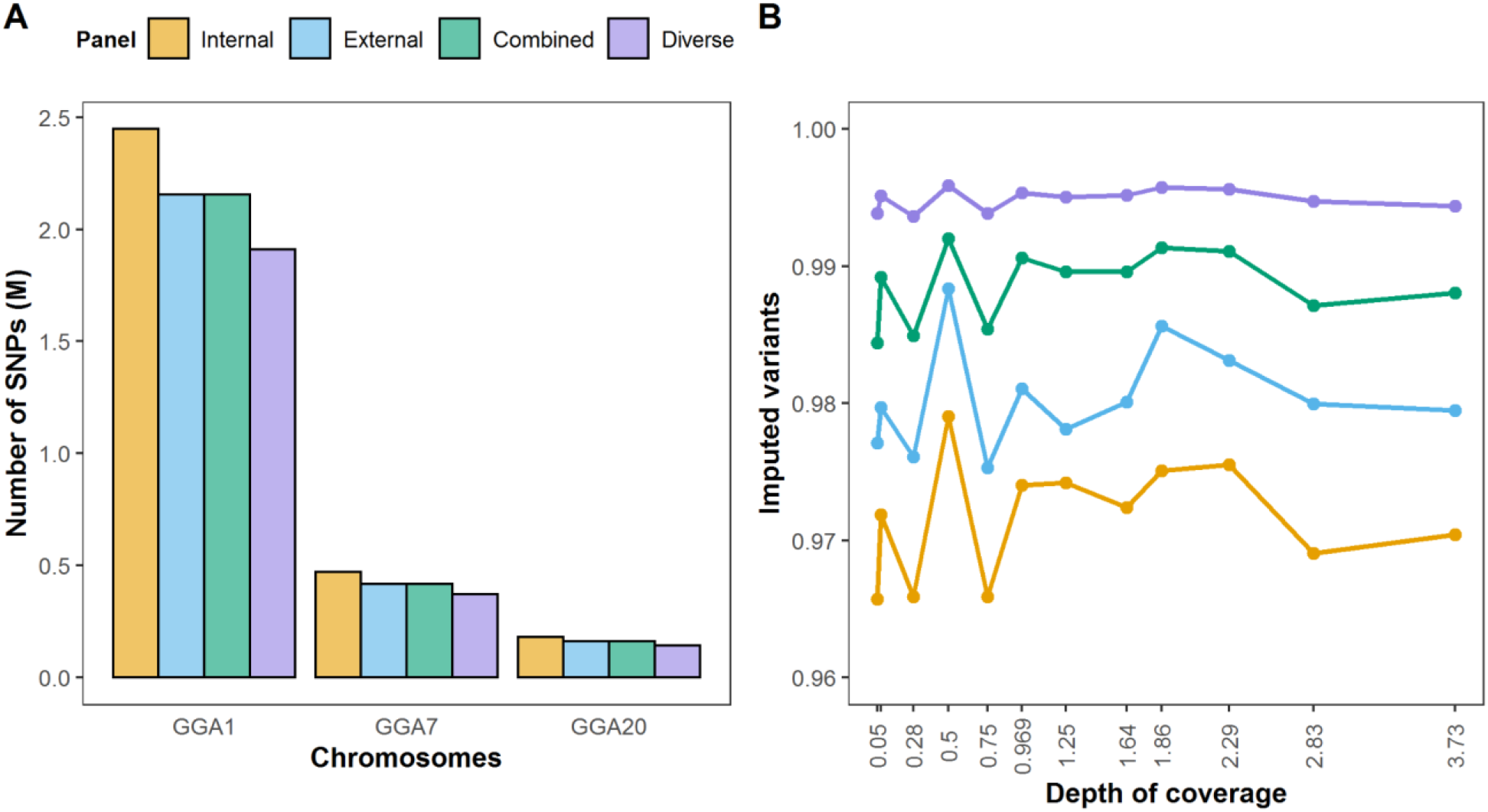
Imputation statistics. (A) Number of SNPs in each reference panel for chromosomes GGA1, GGA7, GGA20. (B) Depth of coverage and proportion of successfully imputed variants of the 12 validation samples for the three chromosomes tested. Capitalised letters refer to panel names: I=internal, E=external, C=combined and D=diverse.

For each imputation scenario, genotype concordance and precision of het. sites were assessed in the 12 validation samples by comparing imputed and true genotypes per individual. Depth of coverage of low-depth samples ranged from 0.05x to 3.5x, and breadth of coverage from 10% to 80%. After performing LOOCV with the four reference panels, average values of genotype concordance for the 12 validation samples exceeded 0.90 for every panel (Fig 3A) and precision of het. sites ranged from 0.78 to 0.91 (Fig 3B). According to Kruskal Wallis tests, the values of concordance (p-value_internal_ > 0.85, p-value_external_ > 0.85, p-value_combined_ > 0.95 and p-value_diverse_ > 0.95) and precision of het. sites (p-value_internal_ > 0.95, p-value_external_ > 0.85, p-value_combined_ > 0.85 and p-value_diverse_ > 0.85) did not differ across chromosomes. However, mean values differed between panels for each chromosome (Fig 3). Concordance values significantly differed when comparing the internal, external and combined panels (Fig 3A). But no differences were detected between the combined and the diverse panels, indicating that no significant increase in imputation accuracy can be achieved in terms of overall concordance by adding more distant individuals. For precision of het. sites, differences were detected for all panels (Fig 3B), including for the combined and the diverse except for GGA20. This suggests that the heterozygous positions are the most sensitive to the imputation process.

**Fig 3.**
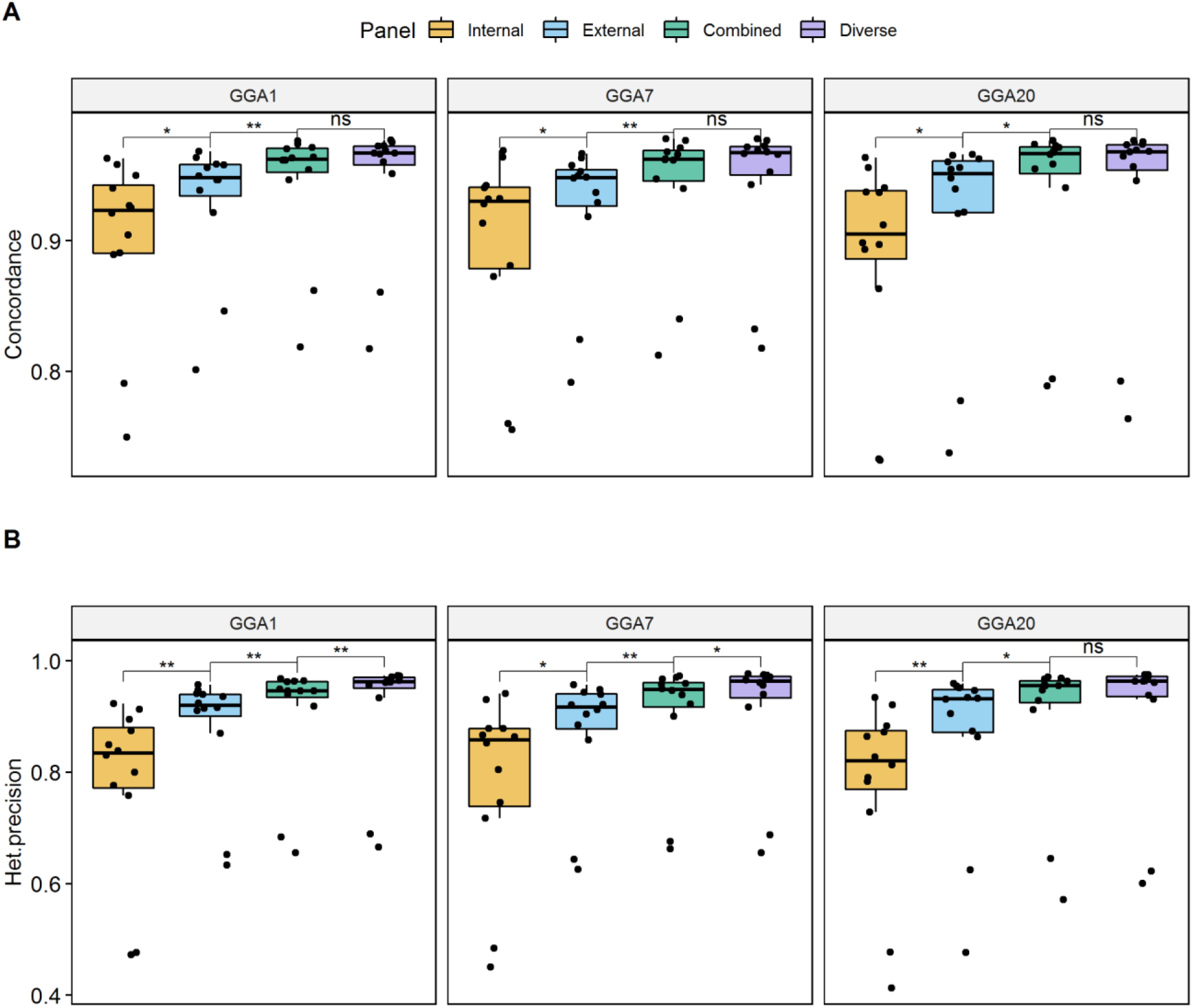
LOOCV test results and comparison of imputation reference panels. (A) Genotype concordance, and (B) precision of heterozygous sites between imputed (low-depth 12 validation samples) and true (internal reference samples) genotypes on chromosomes GGA1, GGA7 and GAA20. Paired T-tests were performed to identify significant differences in means: the following symbols ( “**”, “*”, “ns”) indicate different p-value cut-points (>0.001, 0.001, 0.05).

In an attempt to further assess imputation accuracy, we classified variants according to their MAF in four bins (0-0.05, 0.05-0.1, 0.1-0.3 and >0.3) and calculated precision of het. sites, and the number of correctly imputed variants for the 12 validation samples for GGA1 (Fig 4). The internal panel, while recovering the largest number of variants, was also the panel with the lowest performance in adequately inferring low-frequency variants, especially for the variants with MAF <0.1 (Fig 4A). Although there was no improvement from the external to the combined panel for the smallest MAF bin, a substantial improvement was seen for the rest of the bins. Some significant differences but not as pronounced were also observed from the combined to the diverse. Therefore, the combined panel showed overall the best results with the highest number of correctly imputed variants in all MAF bins (Fig 4B), while maintaining a very similar number of imputed SNPs as the external panel. The diverse panel inferred fewer low-frequency variants, but did so more effectively (Fig 4).

**Fig 4.**
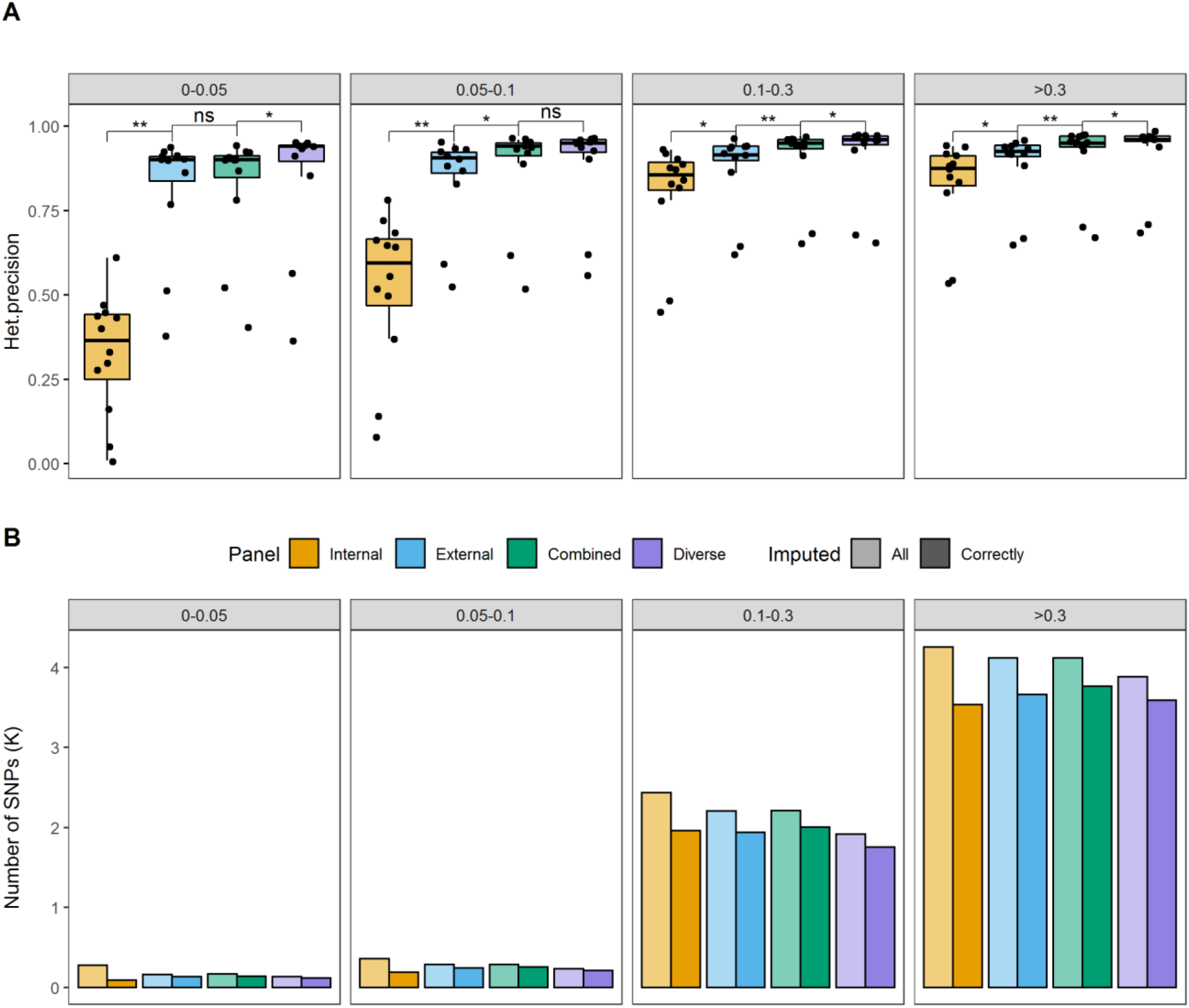
Minor allele frequency variants of LOOCV test. (A) Precision of heterozygous sites and (B) number of imputed low-frequency variants for chromosome one (GGA1) divided into four different bins of minor allele frequency ranges: 0-0.05, 0.05-0.1, 0.1-0.3 and >0.3. The lower bars represent correctly imputed variants, while the bars with greater transparency represent the number of all imputed variants within the respective MAF bin. Variants that coincided between imputed (low-depth 12 validation samples) and true (internal reference samples) genotypes were considered correctly imputed variants. Paired T-tests were performed to identify significant differences in means across panels: the following symbols ( “**”, “*”, “ns”) indicate different p-value cut-points (>0.001, 0.001, 0.05).

Despite the high overall imputation accuracy, the two samples with depths of 0.05x and 0.07x were outliers that did not achieve a sufficiently high concordance (>0.90) and precision (>0.75) with any of the panels and chromosomes (Fig 3). They were thus excluded from the target population, and we refer from now on to 10 validation samples instead of 12.

### Panel choice impact on population genetic inference

#### Number of variants and their allele frequency distribution in the imputed target population

The final number of SNPs recovered from all autosomal chromosomes in the target population with different panels decreased as more distant individuals were included (Fig 5A). This was due to the missing call rate (MCR) filter during the two-step imputation. Using the internal panel, we recovered 11.7 M filtered SNPs in the target population. These were 30% more recovered variants than when using the diverse panel (8.9 M). Most of the excess variants from the internal panel are low-frequency variants that cannot be confidently recovered (Fig 5B), as seen in the less effective imputation of low-frequency variants with the internal panel (Fig 4A). Both Ross and Cobb populations showed extreme allele frequencies (peaks at both ends of the distribution, Fig 5B) revealing a high proportion of fixed or nearly fixed variants in the respective populations. The Ross population had a higher density of low-frequency variants than Cobb (Fig 5C), indicating a higher number of fixed variants than in the Cobb population.

**Fig 5.**
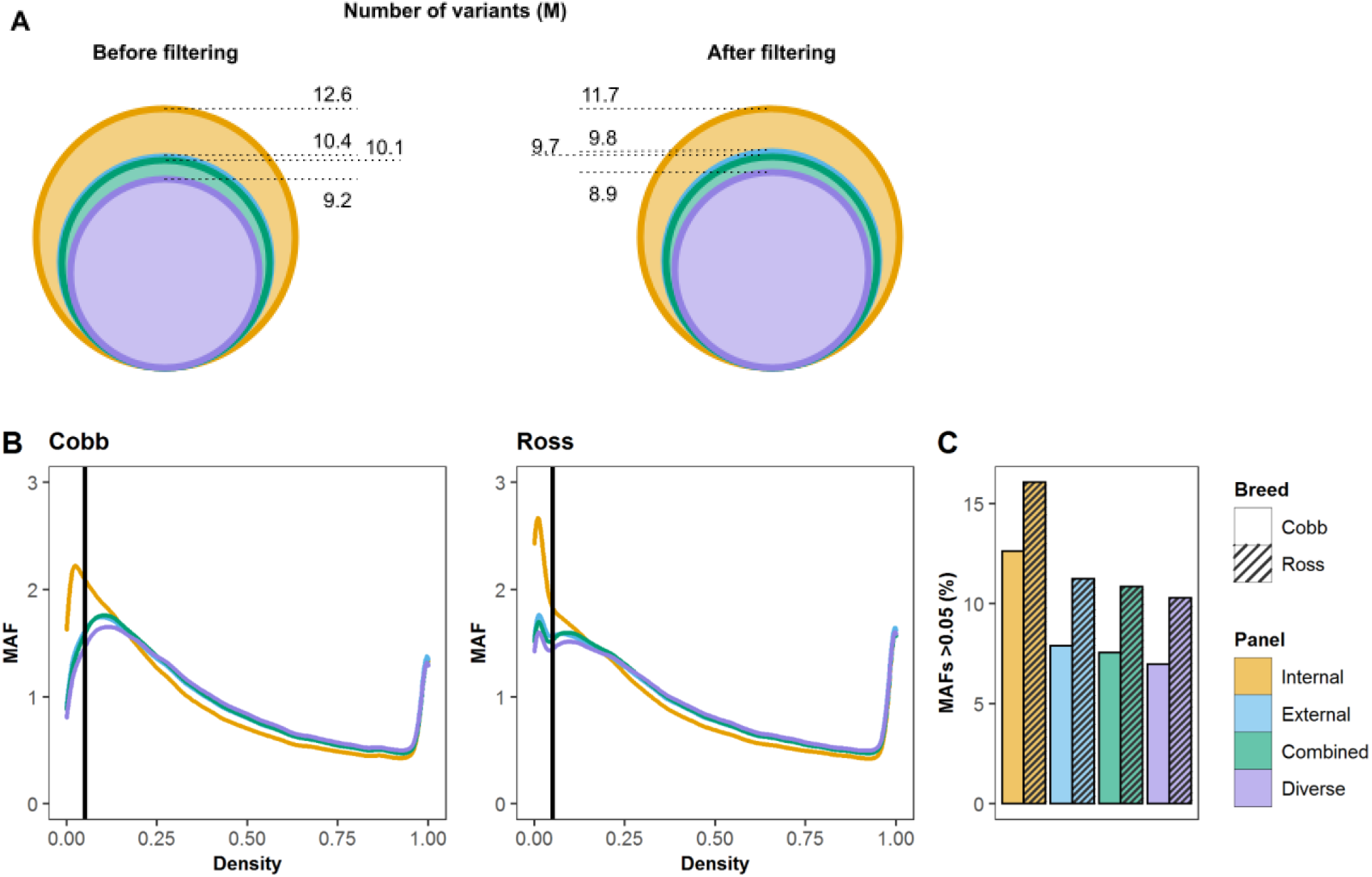
Imputed variants in the target population and their allele frequencies for all autosomal chromosomes. (A) Number of variants in the target population when imputed using the different panels. (B) Allele frequencies of variants imputed in the target population, with a vertical line indicating minor allele frequency (MAF) 0.05, a standard threshold for quality control filtering in genomic datasets. (C) Percentage of variants with a MAF lower than 0.05 by breed for all the panels.

### Population genetic parameters in the target population

In order to explore the effect of panel choice in downstream analyses, we measured five parameters commonly used in population genetics; namely, observed heterozygosity (O.Het), nucleotide diversity (π), fixation index (FST), pairwise distance as measured by 1-identity-by-state (1-IBS) and kinship.

Mean O.Het values differed across all panels for both Cobb and Ross (Fig 6a). The values estimated by imputation tended to increase with panel size and diversity for both breeds. Individual O.Het percentage values displayed a higher variance when imputed with the internal panel and tended to equalise across samples with the external, combined and diverse panels, following the same trend as with the accuracy statistics (Figs 3 and 6A). This high variance displayed by the internal panel might stem from the fewer correctly imputed variants in the internal panel. For the Cobb population, none of the panels reached the heterozygosity values seen with the 4 Cobb individuals (from the high-depth validation samples) (Fig 6A). For Ross, on the contrary, the external and combined panels showed very similar values to the validation samples, while the diverse panel overestimated O.Het values. The very same trend can be seen when comparing imputed and high-depth validation samples (Fig S3). There were some outlier samples (two from Cobb and one from Ross) that presented lower O.Het than the high-depth validation samples (Fig 6A). These samples apparently underwent an incorrect imputation process, but it was not necessarily related to a low mapping depth.

**Fig 6.**
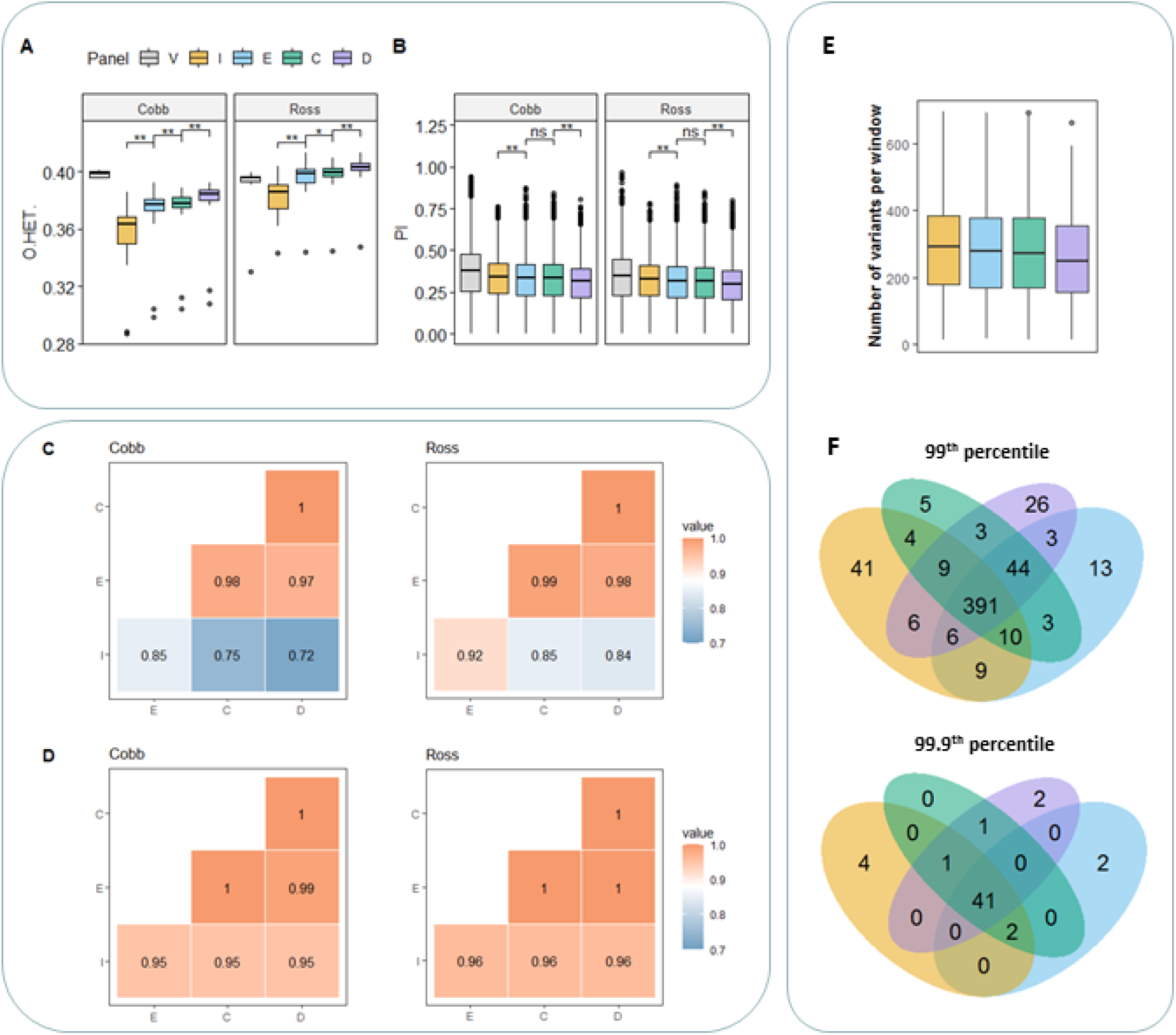
Comparison of the choice of reference panels for imputed target population for all autosomal chromosomes. (A) Observed heterozygosity for the 10 validation samples (true genotypes) and for the imputed target population by breed. Capitalised letters in the legend refer to the following names: I=internal, E=external, C=combined D=diverse and V=Validation samples. (B) Nucleotide diversity of the target population by breed. Paired T-tests were performed to identify significant differences in means: the following symbols (“**”, “*”, “ns”) indicate different p-value cut-points (>0.001, 0.001, 0.05). (C) Kinship and (D) pairwise distance correlation matrices for the target population. Capitalised letters in the x and y axes refer to panel names: I=internal, E=external, C=combined and D=diverse. (E) Boxplot showing number of variants in the common windows of the 99th and 99.9th percentiles from the FST genome scan. (F) Venn diagram depicting overlap of significantly differentiated windows as estimated by FST genome scans between Cobb and Ross populations using the different panels for imputation. Significance thresholds were set at the 99th and 99.9th percentiles.

Nucleotide diversity, on the other side, decreased with increasing panel size and diversity (Fig 6B), which was directly related to the lower number of variants retained in the external, combined and diverse panels compared to the internal. There were significant differences in means except between the external and combined panels for both breeds, most likely because of the similar number of variants both panels share (Fig 5A). When comparing the imputed population with the validation samples, π of imputed samples and of the target population were underestimated for all panels (Figs 6B and S3).

Regarding the population genetic interpretation, both populations were very similar, but the imputation tended to accentuate differences between the two populations (Figs 6 and S4). Within population pairwise distance and kinship values did not vary much according to the panel. For pairwise distance, the diverse panel resulted in larger interindividual distances within breeds (Fig S4). Kinship values were lower when computed with the internal panel, since a larger number of SNPs were retained, in particular, low-frequency variants which are typically unique to one or few individuals thus decreasing kinship (Fig S4). Mantel R tests did not show any significant differences for pairwise distance and kinship matrices, giving the same result for all panel comparisons (Mantel statistic, p-value < 0.001). Correlation values for pairwise distance were very similar and close to 1 (Fig 6D), even for the validation samples (true genotypes) when compared with any panel (Fig S3). For kinship instead, it seemed that the internal panel differed more from the rest (Figs S3 and 6C). In both cases, the 10 validation samples were most correlated with samples imputed with the combined and diverse panels (Fig S3).

Whole-genome mean FST values between Ross and Cobb populations were very similar (internal 0.071, external 0.071, combined 0.072 and diverse 0.072) indicating overall low differentiation between the breeds. When analysing the putative selective sweep regions using as threshold the 99th percentile, 68.2% of the windows coincided across the four panels, but more interestingly, 75.9% of the windows were shared across the external, combined and diverse panels. When we raised the threshold to the 99.9th percentile, 77% of windows were identified by the genome-scans regardless of the choice of panel, indicating that the strongest signals are detected with any panel. Yet, there were some regions that only passed the threshold when imputation was performed with a particular panel (Fig 6F). The combined panel did not show specific sweeps when the percentile was set at 99.9, and it was the panel with the lowest panel-specific regions with the 99th percentile as well, potentially indicating the most robust results, i.e. without panel-specific biases. Surprisingly, the diverse panel detected the most panel-specific sweeps after the internal panel (Fig 6F). On the other side, in terms of density of variants in the common windows, the mean number of variants reduced significantly from the internal to the diverse panel (Fig 6E). This suggests that although in a broad sense the same sweep signals can be detected by all panels, a reduced number of imputed variants might give a smaller chance of detecting causative variants.

## Discussion

Shotgun metagenomic datasets of host-associated microbial communities often contain host DNA that is usually discarded because the amount of data is too low for accurate host genetic analyses. Here, we introduced an effective and accurate approach to recover high-quality host genomes from gut metagenomic data, which can be used to study host population genetic analyses and ultimately contribute to a better understanding of host-microbiota interactions.

Our analyses yielded drastic differences in mapping statistics between caecum samples used to characterise the target population and ileum samples employed to generate the internal reference panel. Although both sample types derived from gut contents, the caecum harbours a very small amount of the host DNA compared to the ileum, because the latter is known to contain fewer bacteria (39), and the higher permeability and a thinner mucus layer of the ileum probably entails higher release of epithelial cells to the lumen (40). Moreover, the low, yet variable, proportion of host DNA retrieved from caecum samples renders sequencing depth adjustment highly unpredictable, as previously reported (7). Notwithstanding, we showed that if a proper reference panel is designed, the low and variable fractions of host DNA recovered from such suboptimal samples, can be used for accurately inferring host genetic features. It must be noted though, that the ratio of host and microbial DNA recovered from chicken caecum and ileum samples can not directly be extrapolated to other host taxa and sample types.

The two-step imputation strategy performed efficiently despite the structural (e.g., study design, animal taxa, reference panel size) differences between our study system and the ones the strategy was originally designed for (23,24). First, we used custom reference panels with less than one hundred individuals, while the two-step strategy was originally tested with large reference panels such as the Human 1000 Genomes (12). Nevertheless, our accuracy values were comparable to the previous results, most likely because the individuals in our target population were closely related, as evidenced by the high kinship values. Second, although we had a similar range of target population sample depths (Hui: from 0.05x to 2x and Homburger: 0.54x to 1.76x), our samples consisted of real low-depth sequence data, instead of downsampled sequencing reads from high-depth samples. Thus, mapping gaps across the reference genome were unevenly distributed. This is evidenced by the large difference between depth (1.8x) and breadth (50%) of coverage (S1 Table), likely hampering accurate computation across the genome. Besides, Hui et al. (2020) documented that the proportion of correctly imputed heterozygous sites started decreasing at 0.5x of depth of coverage, reaching 50% of correctly imputed sites at 0.1x. In our system, >90% of the variants in samples with 0.28-0.5x could be recovered, and accuracy only dropped significantly in samples below 0.1x. Accordingly, we decided to set a mapping depth threshold at 0.28x, but we recommend adjusting it depending on the sample size and quality of the data set, as well as the accuracy needs of each study.

The accuracy of low-frequency variants for all panels except for the internal, which showed much lower values, were comparable to previous works (24), most likely owing to the stringent filtering criteria applied in our study (MCR = 0). But the overall accuracy and the accuracy of heterozygous sites depends heavily on variant frequencies, therefore these comparisons should not be decisive. Finally, unlike humans, avian genomes present macro- and micro-chromosomes and the latter frequently undergo interchromosomal translocations (41). However, it seems that the possible interchromosomal translocations of the target population did not affect imputation, since we did not find any significant differences in accuracy between chromosomes, revealing that the strategy worked equally well for large, mid-sized and small chromosomes with potentially different linkage patterns.

### 4.1 Effect of reference panel on accuracy statistics

Reference panel design depends on data availability as well as computational capacity. It is a common strategy for imputation of inbred populations to resequence a subset of samples with higher resolution in order to optimise imputation performance (35). Based on previous works, we estimated that 12 individuals out of 100 would be sufficient to represent the genetic diversity of the population. For instance, previous chicken studies deep-sequenced 25 individuals to impute approximately 450 chickens genotyped with 600-K SNP arrays (~5% of sample size) (20,42).

In terms of panels SNP density, we decided to genotype variants that did appear in our target population rather than calling for specific variants in the rest of the breeds that composed the reference panels. Thereby, we aimed at reducing the noise that the excess of variant density could cause in the imputation process. Nevertheless, as the genetic distance between the selected and our breeds is very small (43), we expected them to share many variants, as we evidenced with preliminary analyses using GGA1 where 72% of variants identified by genotyping or by calling overlapped in the external panel (Fig S2).

The internal panel resulted in a larger variance across samples. SNPs with low MAF had the lowest accuracy when imputed with the internal panel. Moreover, incorrectly imputed low-frequency variants can be easily overcome if a strict MAF filter is applied for downstream analysis. Another possible option is to sequence more individuals of the target population to increase the reference panel size. Hence, despite the internal panel only representing a small subset of the target population, and showing lower imputation values than in the external, combined, and diverse panels, for scientists without access to external reference samples, this approach is equally useful as overall imputation accuracy was higher than 90% and biological differences were still visible. In this sense, host resequencing of a small subset of the target population might represent a cost-efficient option, especially for researchers working with non-model organisms and inbred populations. Thus, our approach could be useful, for example, to study genome features of endangered populations relying on faecal samples recovered from the environment.

Our results showed that the combined panel performed better in terms of overall accuracy, and specifically of minor allele frequency variants, than the internal and the external panels alone. Despite the fact that the external and combined panels had the same number of SNPs, including a subset of individuals from the target population was beneficial. Many studies already mentioned an improvement for the combined option (44,45). Lastly, the diverse panel showed the highest values of concordance and het. sites precision, most probably because of the lower number of SNPs recovered, especially low-frequency variants, which generally yielded lower imputation accuracies. In terms of imputation of low-frequency variants, the combined panel outperformed the diverse one, i.e. it correctly imputed a larger number of variants and tended to improve the precision of het. sites in some MAF bins. A recent large-scale study performed in a Chinese population showed that a population-specific reference panel worked the best compared to European reference panels such as 1000G (21). Imputation was greatly improved when the reference panel contained a fraction of an extra diverse sample, but they obtained a different pattern when the panel size was fixed (21). Thus, taking into consideration our and previous results on selection of imputation panels, it can be concluded that increasing panel size and diversity improves imputation, but a balance has to be found in the composition of the panel. The distance between the panel and the target population has to be taken into account.

### 4.2 Effect of reference panel on biological inference

Besides crude imputation accuracy statistics, we evaluated the impact of the panels on downstream population genetic parameters and their biological interpretation. As imputation accuracies were generally high with our applied pipeline and the stringent filtering approach, we expected population genetic inferences to follow similarly.

Although overall results were in agreement, all the tested parameters showed slight trends according to the used reference panel. O.Het, pairwise distance and kinship values increased while mean FST and π values decreased with panel size and diversity (Figs 6 and S4). Such biases were related to the composition of the panels and the associated number and distribution of recovered SNPs.

Imputation worked slightly differently for the two breeds, as Ross population estimations were closer to the true values than for the Cobb population. Thus, accentuating the distance between both breeds. This is most likely due to a smaller representation of Cobb individuals in the reference panels, i.e. 5 Cobb and 7 Ross samples constituted the internal reference panel. Secondly, there were some samples that were incorrectly imputed because of their low O.Het values (Fig 6A). We do not know if there are individuals with lower heterozygosity in our Cobb and Ross populations. For instance, there was a Ross individual from the high-depth validation samples with lower O.Het. Chickens came from two different hatcheries, which might be the reason why some individuals might have slightly different genetic features. We may have under-represented one of the origins in the internal reference samples. Thus, it is necessary to be more cautious for the interpretation of individual genomes. Nevertheless, results appeared to be robust and similar across panels at the population level. The genome scans yielded overall very consistent results with major differentiation signals identified by any of the imputed datasets, likely indicative of a true selection signature between both breeds. However, downstream analyses such genome scans and GWAS must be performed with caution since this method is sensitive to low-frequency variants quality.

Both breeds exhibited extreme minor allele frequencies, indicating that the genetic drift due to selection in a closed breeding population has a notable effect. Domestication and breeding history are the two major processes that shape haplotype structure (31,46). Cobb and Ross, together with other commercial breeds, have much smaller effective population size than other chickens (47). Broiler breeding methods are described as a pyramid strategy, in which pure, inbred lines are crossed, then F1 individuals are crossed between each other. In some cases, even a second or a third cross is performed in F2 and F3 generations before raising them for meat (48). Therefore, broilers are highly related populations, but at the same time present high heterozygosity values. Heterozygosity of our studied breeds were much higher O.Het than of local populations (49), but similar to other broiler breeds (46). Similarly, nucleotide diversity and mean fixation index values were comparable to those previously reported (31).

## Conclusions

Our results show that the two-step imputation implemented in this study can be used to successfully reconstruct genotypes and study population genetic properties of hosts from suboptimal metagenomic samples. The comparison among reference panels also demonstrated that this method is versatile and flexible. This approach could be used in many contexts and exploit different data sources to address a variety of research questions. This includes the possibility of mining published metagenomic data sets to recover discarded host DNA sequences. In our particular case, the reconstructed genotypes will be employed in the H2020 project HoloFood to detect interactions with microbial metagenomic features, and thus implement a hologenomic approach to improve animal production (50). Because ‘host-contamination’ should no longer be considered a problem, but an opportunity.

## Acknowledgments

We would like to thank the partners that were involved in the design and execution of the animal trials, specially our colleagues Joan Tarradas, Nuria Tous and Enric Esteve from IRTA.

## Supporting information

**S1 Fig. Alignment results before and after changing seed length from 19 to 25.** Captures from multiple regions of the GGA1 visualized with Geneious.

**S1 Table. Mapping depth and breadth results before and after changing seed length from 19 to 25.**

**S2 Table. Individual mapping depth and breadth values of the target population.**

**S2 Fig. Venn diagram of shared variants between the internal reference samples and the variant called 40 broilers of the external panel for GGA1.**

**S3 Fig. Comparison of the choice of the reference panel for the imputed 10 validation samples.** (A) Observed heterozygosity, (B) nucleotide diversity and correlation plots for (C) pairwise distance and (D) kinship were measured to compare imputed and true genotypes of the validation samples.

**S4 Fig. Pairwise distance and kinship heatmap matrices for each of the panels.**

## Notes

### Competing Interest Statement

The authors have declared no competing interest.

